# Dietary restriction fails to extend life in stressful environments

**DOI:** 10.1101/2022.10.17.512576

**Authors:** Felix Zajitschek, Susanne R.K. Zajitschek, Ana C.O. Vasconcelos, Russell Bonduriansky

## Abstract

Moderate dietary restriction often prolongs life in laboratory animals, and this response has been interpreted as an adaptive strategy that promotes survival during famine. However, dietary restriction can also increase frailty, and it therefore remains unclear whether restricted diets prolong life under stressful conditions like those experienced by wild animals. We manipulated adult dietary protein of *Drosophila melanogaster* across a gradient of ambient temperature. We found that protein restriction increased longevity of both sexes at benign ambient temperatures (25-27°C), but failed to extend or even reduced longevity of flies maintained in cold (21-23°C) or hot (29°C) conditions. Protein restriction also generally reduced reproductive performance, and did not consistently enhance performance of F1, F2 or F3 descendants. Our results challenge the long-held idea that extended longevity of diet-restricted laboratory animals represents an adaptive survival strategy in natural populations, and suggest instead that this response is an artefact of benign laboratory conditions.

## INTRODUCTION

It has been known for decades that moderate dietary restriction (DR) extends longevity in a wide variety of experimental animals (Fontana *et al*. 2010). Many studies have investigated the specific diet components (especially calories, protein, or specific amino acids) whose restriction can induce this response (Piper & Partridge 2007; Lee *et al*. 2008; Grandison *et al*. 2009; Lee 2015; Soultoukis & Partridge 2016), and the physiological mechanisms involved (Zanco *et al*. 2021; Green *et al*. 2022). While DR is seen as a potential means to extend human lifespan and healthspan (Heilbronn & Ravussin 2003; Maegawa *et al*. 2017; Pifferi & Aujard 2019), the life-extending effects of DR are also of considerable interest from an evolutionary perspective. However, the evolutionary interpretation of these effects remains controversial (Kirkwood & Rose 1991; Kirkwood & Shanley 2005; Adler & Bonduriansky 2014; Moatt *et al*. 2020; Piper *et al*. 2022).

An influential evolutionary hypothesis posits that the physiological responses that enhance longevity when nutrient intake is restricted represent an evolved strategy that functions to enable animals to survive periods of famine (Holiday 1989; Kirkwood & Rose 1991; Kirkwood & Shanley 2005). According to this “adaptive resource reallocation” hypothesis, DR induces a reallocation of metabolic resources from reproduction to somatic maintenance, and an up-regulation of cellular recycling and repair processes such as apoptosis, because these responses reduce mortality and thereby increase the probability of surviving until resources become abundant again and reproduction can resume. This hypothesis is supported by evidence that *Drosophila melanogaster* females are able to resume full reproduction after a period of DR (Sultanova *et al*. 2021), although increased nutrient abundance following a period of DR can result in elevated mortality that could negate the reproductive gains (McCracken *et al*. 2020).

Responses to DR have been likened to seasonal diapause, when reproduction ceases and specialised survival mechanisms are activated (Regan *et al*. 2020). However, a key assumption of the adaptive resource reallocation hypothesis is that the strategy of foregoing reproduction while resources are scarce can pay off in wild insects and other animals *during the breeding season*. This assumption is problematic in light of the ecology of natural populations and the physiological effects of DR. Although estimates of mortality rate in wild insects are scarce, the available data suggest that reproductively active adult flies experience a mortality rate of ∼ 10% per day (Bonduriansky & Brassil 2002; Kawasaki *et al*. 2008; Mautz *et al*. 2019a), and estimates from many other insects are similarly high (Zajitschek & Bonduriansky 2014; Zajitschek *et al*. 2019). Such a high risk of death means that any delay in reproduction is likely to be very costly for most insects and other small-bodied animals. This problem could be overcome if physiological responses to DR substantially reduce mortality risk in the wild, but there is little evidence to suggest such an effect. Mair (2005) found that DR enhanced the ability of *Drosophila melanogaster* females (but not males) to resist starvation, but reduced females’ ability to resist desiccation; no effect was observed on resistance to cold, heat, or oxidative stress (paraquat). Other studies have reported increased frailty in nutrient-restricted animals subjected to stressful conditions, including a reduced capacity to mount an immune response against pathogens, reduced ability to heal wounds, and reduced cold tolerance (reviewed in Adler & Bonduriansky 2014). For example, in *Drosophila ananassae*, dietary protein enhanced desiccation and heat-shock resistance (Sisodia & Singh 2012) while, in *Drosophila melanogaster*, wing clipping reduced longevity of protein-restricted flies but not of fully fed flies (Ghimire & Kim 2015). The ability to tolerate environmental challenges such as thermal stress and wounding is relatively unimportant in the typically benign conditions of the laboratory, but is likely to have a considerable impact on survival of wild animals living in natural environments.

Furthermore, in insects and other small-bodied animals, many physiological processes are tightly coupled to ambient temperature (Sestini *et al*. 1991; Keil *et al*. 2015; Mołoń *et al*. 2020). Consequently, physiological responses (such as reduced metabolic rate) and behavioural strategies (such as reduced activity) that can be deployed to reduce mortality risk during cold-season diapause might be limited or impossible during the breeding season, regardless of nutrition. Moreover, protein restriction induces increased activity levels in *D. melanogaster* (Ghimire & Kim 2015; Krittika & Yadav 2020), and this behavioural response could elevate risk of predation for protein-restricted animals in the wild.

Assessing the adaptive resource reallocation hypothesis therefore requires verifying the key premise that nutrient limitation can increase survival of non-diapausing animals not only under benign conditions but also in challenging, stressful environments. If nutrient limitation fails to reduce (or even increases) mortality risk under stress, then life-extension under DR is likely to be an artefact of laboratory conditions rather than an evolved strategy that enhances fitness in natural populations.

Much of the evidence on the lifespan-extending effects of DR comes from experiments with flies and other insects, and a general finding across many insect species is that lifespan is extended when protein intake is reduced at the adult stage (Maklakov *et al*. 2008; Adler *et al*. 2013; Lee 2015; Regan *et al*. 2020; Zanco *et al*. 2021), and perhaps also during development (Runagall-McNaull *et al*. 2015). In the most important model insect system for DR research, *Drosophila melanogaster*, it is well-established that protein-restricted flies typically outlive fully fed flies, but protein-restricted females tend to be less fecund (Lee *et al*. 2008; Lee 2015; Simpson *et al*. 2017; Sultanova *et al*. 2021). However, the vast majority of DR experiments on *D. melanogaster* and other insects have been carried out under benign laboratory conditions, including a stable ambient temperature to which the flies are well-adapted (25-27°C), few pathogens and parasites, and no predators. Only a few studies have investigated effects of DR on longevity in more challenging environments, and these studies have reported strongly context-dependent effects of protein consumption that cast doubt on the idea that protein-restricted animals typically achieve extended longevity in natural environments. Burger *et al*. (2006) found that diluted diets reduced mortality from bacterial infection but also reduced oxidative stress resistance and cold-tolerance in *D. melanogaster* females. Savola *et al*. (2021) found that restricted access to dietary protein increased mortality of *D. melanogaster* females that had been infected with a bacterial pathogen, whereas survival of injured females was not affected by dietary protein. Mautz *et al*. (2019a) manipulated access to protein in both captive and wild cohorts of male antler flies (*Protopiophila litigata*), and found that protein-supplemented adult males exhibited reduced longevity in the lab but not in the wild, although this protein × environment interaction was supported in only one of the two years of the study.

Here, we asked whether DR (protein restriction) enhances longevity of *D. melanogaster* females and males not only in benign environments but also under thermal stress. Temperature fluctuations are experienced by all natural populations, and their effects are likely to pose especially severe challenges to small-bodied, ectothermic animals such as insects. Ambient temperature affects the expression of a range of physiological and life-history traits in *D. melanogaster* (Sestini *et al*. 1991; Mołoń *et al*. 2020), and exposure to both cold and hot temperature extremes can be stressful (Arias *et al*. 2012; Mockett & Matsumoto 2014; Klepsatel *et al*. 2016). The potential for temperature stress to modulate the effects of DR on lifespan is therefore especially relevant to understanding how DR might affect fitness in wild flies and other wild animals. We manipulated adult dietary protein (40%, 100% and 150% of the normal yeast concentration, with other diet components held constant) across a gradient of ambient temperature, including below-normal temperature that is likely to induce cold stress (21°C, 23°C), normal rearing temperature to which the flies are likely to be well adapted (25°C, 27°C), and above-normal temperature that is likely to induce heat stress (29°C). Given the potential for trade-offs between survival and reproduction in *Drosophila* (Flatt 2011), we quantified both longevity and reproductive performance (female fecundity, male mating success) to determine whether negative effects of environmental conditions on longevity might be offset by positive effects on reproduction. Likewise, we examined effects of diet and temperature treatments on performance of the offspring, grand-offspring and great-grand-offspring of exposed flies to determine whether treatment effects on exposed individuals might be offset by effects on their descendants (see Mautz *et al*. 2019b).

## METHODS

Experimental flies were sourced from the outbred Dahomey population. This population was started in 1970 from founders caught in the wild in Dahomey (now Benin), West Africa, and has been maintained in population cages, containing several thousand individual males and females, with overlapping generations. Females in population cages were given the opportunity to lay eggs in vials containing standard sugar-yeast (1.0 SY) diet. Eggs were collected and distributed to 100 new 1.0 SY food vials (10 ml food per vial; 50 eggs per vial). Offspring were collected and allowed to mate with offspring of a different vial for the first 48 hours of adult life (20 males and 20 females from each vial, resulting in 40 adults per combined vial). At age 3 days, males were removed, females were transferred into new vials (20 per vial) and allowed to lay eggs for five hours. Egg number per vial was trimmed to 60-70. Flies were reared for another generation, using the same protocols, and the eclosing adults of the third generation were used as focal experimental subjects. Males and females were given the opportunity to mate for 48 hours after eclosion, before the sexes were separated and flies were distributed in treatment groups.

We created three different diets by manipulating the protein content and therefore the protein to carbohydrate ratio (protein content in gram yeast per litre diet: 40 = low, 100 = standard, 150 = rich; Table S1), while maintaining the amount of sugar in each diet (at 50 g / litre diet). The effects of each of the three diets were tested at each of five temperatures (21, 23, 25, 27, 29°C), resulting in 15 treatment groups. The experiment was performed in a controlled temperature and humidity room, at 21°C and 60% relative humidity, with a 12:12 hours light:dark cycle. Treatment temperatures above 21°C were established with heat mats, regulated by digital thermostats, placed in clear plastic containers (52 × 35 × 28 cm). Vials in the 21°C treatment were place in an identical container, containing no heat mat. Vials were placed upright in cardboard trays (on plastic frames, 10 cm above heating mats), on top of a piece of carboard with the same area as the tray (0.5 cm thick in total) to avoid differences in temperature due to potentially localized differences in heat produced by the heating mat (temperatures at 10 cm above heating mats were tested at different locations in each temperature container and not found to be different, prior to the experiment). Vials were randomly allocated within trays every time flies were flipped to new vials. Containers were fitted with lids, leaving a 2 cm wide space open at one short side of the container to allow for air exchange. We monitored temperature inside the boxes with individual temperature loggers, placed at the height of the food surface in experimental vials. Temperatures in the highest temperature treatment never exceeded the set treatment temperature by more than 0.4°C (mean, variance, minimum, maximum for temperature: ambient: 20.58°C, 0.09, 20.20°C, 22.30°C; 21°C container: 20.69°C, 0.06, 20.21°C, 22.15°C; 23°C container: 23.19°C, 0.13, 22.84°C, 23.66°C; 25°C container: ; 25°C container: 24.83°C, 0.11, 23.8°C, 25.42°C; 27°C container: 27.14°C, 0.02, 26.40°C, 27.70°C; 29°C container: 28.58°C, 0.29, 26.6°C, 29.4°C).

For survival estimates, each vial was populated with ten individuals, with ten replicate vials per sex per treatment combination (see Table S2 for more details).

Live flies were transferred to new food vials and the number of dead flies per sex was recorded every Monday, Wednesday, and Friday. Female fecundity was estimated by collecting and counting eggs laid by females that were used to measure survival, over 18 hours, once a week (Wednesday to Thursday), for the first five weeks of the experiment.

### Parental effects and male mating behaviour

To measure male mating behaviour and test for parental temperature and diet treatment effects on offspring, we established additional vials per treatment and sex (6 vials with males, 3 vials with females), containing 20 individuals (male mating behaviour F0 (MMB), paternal effect F0 (PE) and maternal effect F0 (ME)). Males and females experienced the same treatment as the flies used to measure survival, except females were collected as virgins and their fecundity was not measured.

To test for paternal effects, we mated PE males at age 16 days with 5 days old standard virgin females which originated from the same Dahomey population and were bred for 3 generations following the same protocol as used for the focal experimental flies (F0 survival, PE, and ME flies). For this, the 20 PE males per vial were paired with 20 virgin females for 24 hours. Sexes were separated and females were allowed to lay eggs for 24 hours on standard 1.0 SY food vials (7 ml food per vial). To test maternal effects, we mated virgin ME females at age 16 days to 5 days old standard males, originating from the same batch of flies as the standard virgin females that were mated to PE males. Standard males had the opportunity to mate for the first 48 hours of adult life. Similar to PE matings, 20 ME females per vial were paired with 20 standard males for 24 hours. Sexes were then separated and ME females were allowed to lay eggs for 24 hours on standard 1.0 SY food vials (7 ml food per vial). For PE and ME, three vials with eggs were collected and trimmed to 60-70 eggs per vial.

Eclosing F1 offspring were pooled across the three replicate vials within treatment, sexes were separated, and two vials (with 15 individuals each) per sex for each parental sex-specific treatment were established on standard 1.0SY diet, to measure survival and female fecundity, following protocols used for F0. To test male mating behaviour, an additional two vials per sex-specific parental treatment were populated with 10 males each. All F1 offspring were kept in a temperature chamber, set to 25°C, and 60% relative humidity, with a 12:12 hours light:dark cycle.

To investigate potential effects of F0 diet and temperature treatments on grand-offspring and great-grand-offspring, we obtained F2 and F3 flies from eggs laid by F1 and F2 females at age 10 days, respectively. Transgenerational effects can wane over generations (Bonduriansky 2021). To maximize potential to detect effects of F0 diet and temperature treatments on F2 and F3 descendants, we therefore maintained F2 and F3 adult flies at a high ambient temperature (29°C) to gauge their capacity to cope with heat-stress. F2 and F3 flies were maintained on standard 1.0SY diet, and in two replicate vials (with 15 individuals each) per sex for each parental F0 sex-specific treatment combination. For practical reasons, F2 and F3 descendants were obtained only from a subset of F0 sex-specific treatment combinations (21°C, 25°C, and 29°C, and restricted (0.4SY) and rich (1.5SY) diets, resulting in 6 grand-parental treatments and 6 great-grand-parental treatments), and we only tested for effects of F0 diet and temperature treatments on male mating performance.

Male mating performance was tested for 14 males per treatment, 7 sourced from each of two replicate vials, in the parental (F0), offspring (F1), grand-offspring (F2) and great-grand-offspring (F3) generations. First, standard 5 days old virgin females, maintained in groups of 20 females until the behavioural assay, were aspirated into behavioural vials (one female per vial). Vials contained 5 ml of 0.4SY food and were closed with a white foam plug, leaving a 5 cm high (∼20 ml) mating arena. Treatment males were then aspirated and paired 1 on 1 with females. Pairs were randomly placed in the behaviour observation trays. We recorded time until mating (latency to mate), mating duration, and whether a mating took place (mating success). Observations were performed for 3 hours, starting at 9 am, in a controlled temperature room at 25°C.

### Statistical analysis

We analysed survival of F0 flies using generalized additive models (gam, R package mgcv), since mixed effects Cox proportional hazards models (R package coxme) showed violations of the assumption of proportionality of hazards (tested with function cox.zph in package survival). In two separate model sets for males and for females, we compared three models for hazard that either included diet-specific smooth terms for temperature and diet-specific intercepts, diet-specific intercepts with one common smooth term for temperature, or one common smooth term for temperature only. In all models, we included individual vial as a random effect. We used AIC to compare models within sex (R package MuMIn). If a model that contained diet was supported most, we further tested the effect of diet at each temperature with logrank tests and Benjamini & Hochberg correction for multiple comparisons (package survminer). Sex-specific parental effects on survival on male and female F1 flies were tested with sex-specific mixed effects Cox proportional hazards models (R package coxme), since the proportionality of hazards assumption, as tested with function cox.zph in package survival, was supported.

We estimated total fecundity for each vial as total sum of weekly egg counts in the first 5 weeks after establishing the vial. We analysed the square-root of fecundity values in robust general linear models (function glmrob in R package robustbase), as the diagnostics of general linear models showed problematic patterns at both sides of the observed range of data. Temperature was modelled as a cubic B-spline (function bs() in package splines) and was tested in interaction with diet (glm with the same model specifications yielded qualitatively similar results). Diet was included as a numerical variable, corresponding to its protein content, in linear and quadratic form. We tested the effects of model terms by first removing the most complex term, in this case the interaction between temperature and the quadratic term of diet, and comparing it to the full model, using Wald tests.

We analysed male latency to mate in additive mixed models (function gamm4 in R package gamm4) and general mixed linear models (function lmer in R package lme4). Where necessary to meet model assumptions, we compared models with more suitable error distributions and link functions, and models with a transformed response variable.

## RESULTS

### Survival

The most complex model for F0 female survival, which contained diet-specific smooth terms, was supported most, and provided a substantially better fit than the second-best model (Table S4). Survival of females on restricted diet differed from survival of females on standard or rich diets, except in one temperature treatment (25°C; Table S5). Differences were most pronounced under temperatures 21, 23, and 27°C: at 21 and 23°C, females on a restricted diet exhibited drastically reduced survival (resulting from high mortality from around age 20 days); conversely, at 27°C, females on a restricted diet had increased survival (Figure 1). Female survival differed between standard and rich diet groups only at 25 and 29°C, with a moderate survival advantage of females on standard diet, manifested in late life at 25°C and in mid-life at 29°C.

**Figure 1.**
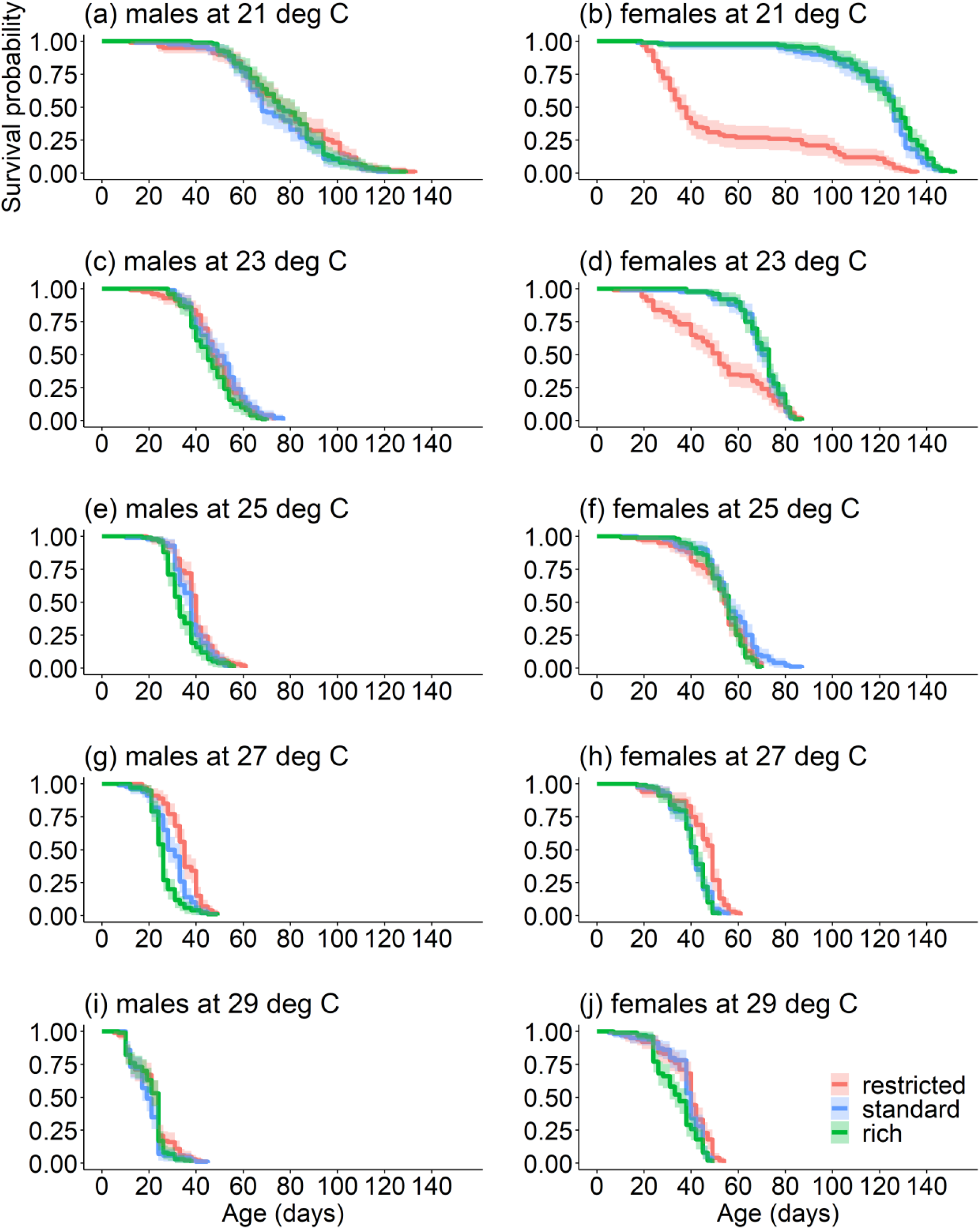
Survival curves of F0 male and female flies. Each panel shows Kaplan-Meier survival curves for sex-specific temperature treatment groups. Separate curves depict survivorship of different diet treatment groups. Shaded areas represent 95% confidence bands. Abbreviations: deg C: degrees Celsius.

Compared to females, male survival in the parental (F0) generation was not as strongly affected by diet (Tables S4, S5). Males on lower protein diets showed a robust survival advantage at 25 and 27°C, but dietary protein did not affect male survival at other temperatures (Figure 1).

Survival of offspring (F1) was not affected by either temperature or diet experienced by male or female parents (Tables S6, S7; model simplification did not change this conclusion).

### Fecundity

The most complex model of F0 fecundity that includes a cubic B-spline of temperature, diet squared, and their interaction provided the best fit (compared to a reduced model without the interaction term diet^2^ × temperature: change in robust quasi-deviance = 40.13, df = 3, p < 0.001). Fecundity of F0 females on the restricted diet was substantially lower than fecundity of females on the standard and rich diets at all temperatures except 29°C (Table S8; Figure 2). For females on the standard and especially the rich diet, fecundity tended to decline with temperature, reaching levels similar to the restricted diet at 29°C (Table S8; Figure 2).

**Figure 2.**
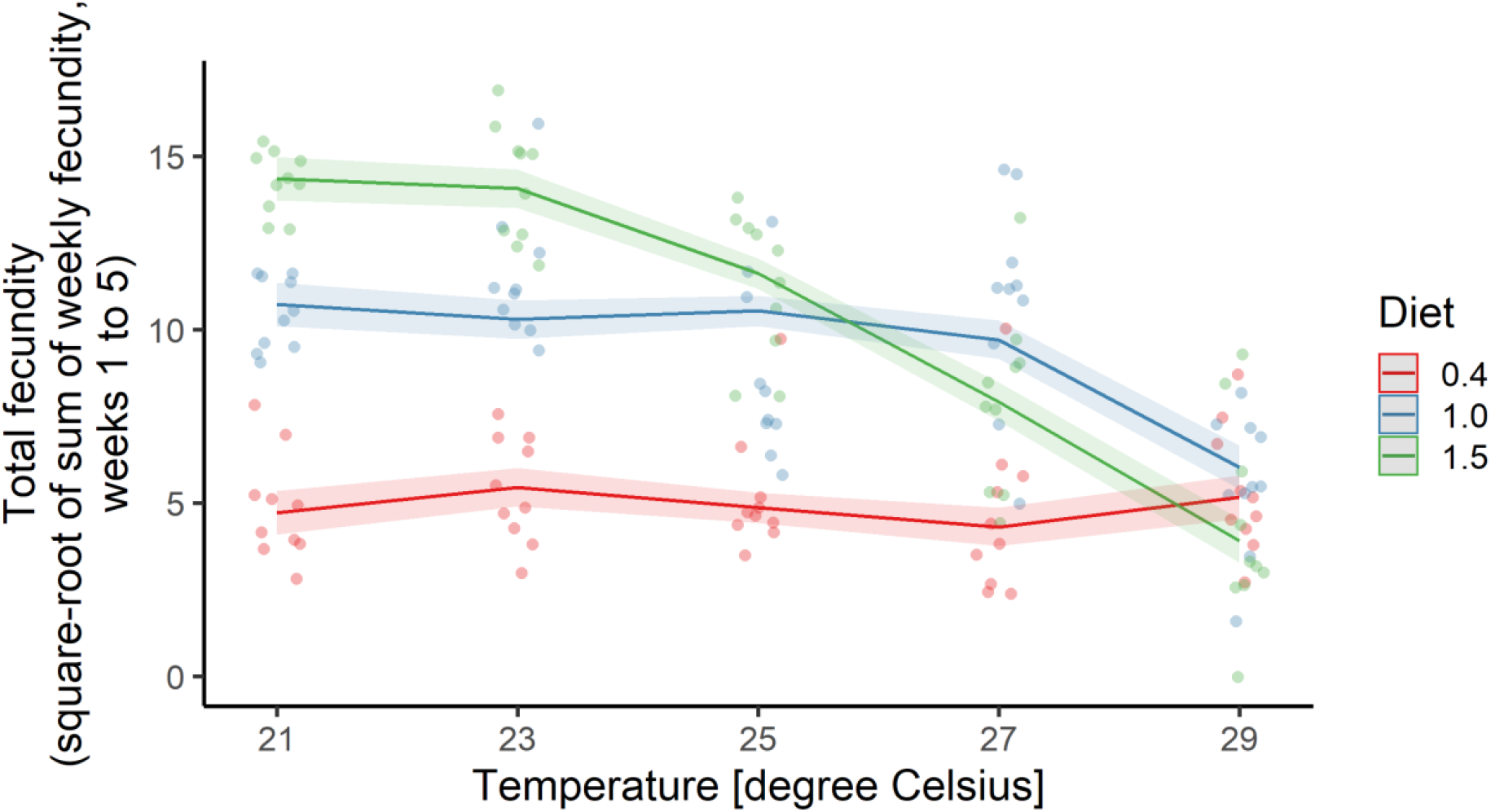
Female fecundity in generation F0. Shaded areas represent 95% confidence bands.

Fecundity of F1 females was also predicted best by the most complex model (compared to a reduced model without the interaction term parental sex effect × temperature: change in robust quasi deviance = 52.28, df = 10, p < 0.001; vs model without parental sex effect × diet: change in deviance = 40.87, df = 7, p < 0.001; model without diet^2^ × temperature: change in deviance = 53.00, df = 9, p < 0.001). For low (21 and 23°C) and high (29°C) maternal temperature treatments, daughters of standard-diet and rich-diet mothers laid more eggs than daughters of restricted-diet mothers, but this pattern was reversed at intermediate maternal temperature treatments (25 and 27°C) (Table S9; Figure 3). For 21, 23 and 25°C paternal temperature treatments, daughters of rich-diet fathers tended to lay fewer eggs than daughters of restricted-diet fathers, but there was no clear effect of paternal diet on daughters’ fecundity when fathers were maintained at 27 or 29°C. Overall, daughters’ fecundity tended to decline with paternal treatment temperature (Table S9; Figure 3).

**Figure 3.**
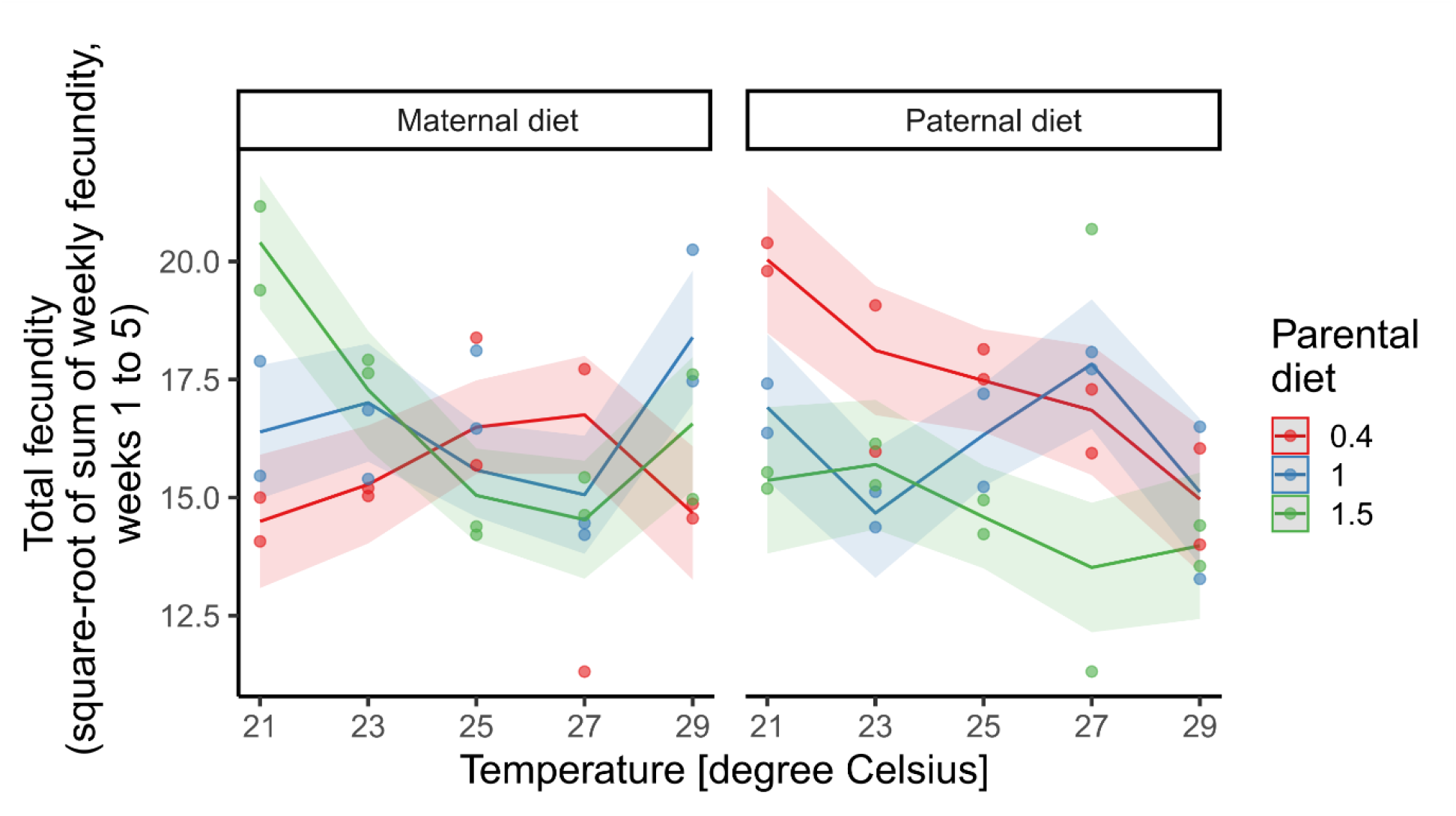
Female fecundity in generation F1. Shaded areas represent 95% confidence bands.

### Male mating behaviour

Latency to mate of males in the parental (F0) generation increased linearly with treatment temperature, independently of diet (Figure 4; Table S10; Temperature-dependent smooth model, approximate significance of temperature smooth term: edf = 1, F = 14.29, p < 0.001). There were no effects of temperature or diet treatments on F0 males’ mating duration or mating success (Tables S11, S12).

**Figure 4.**
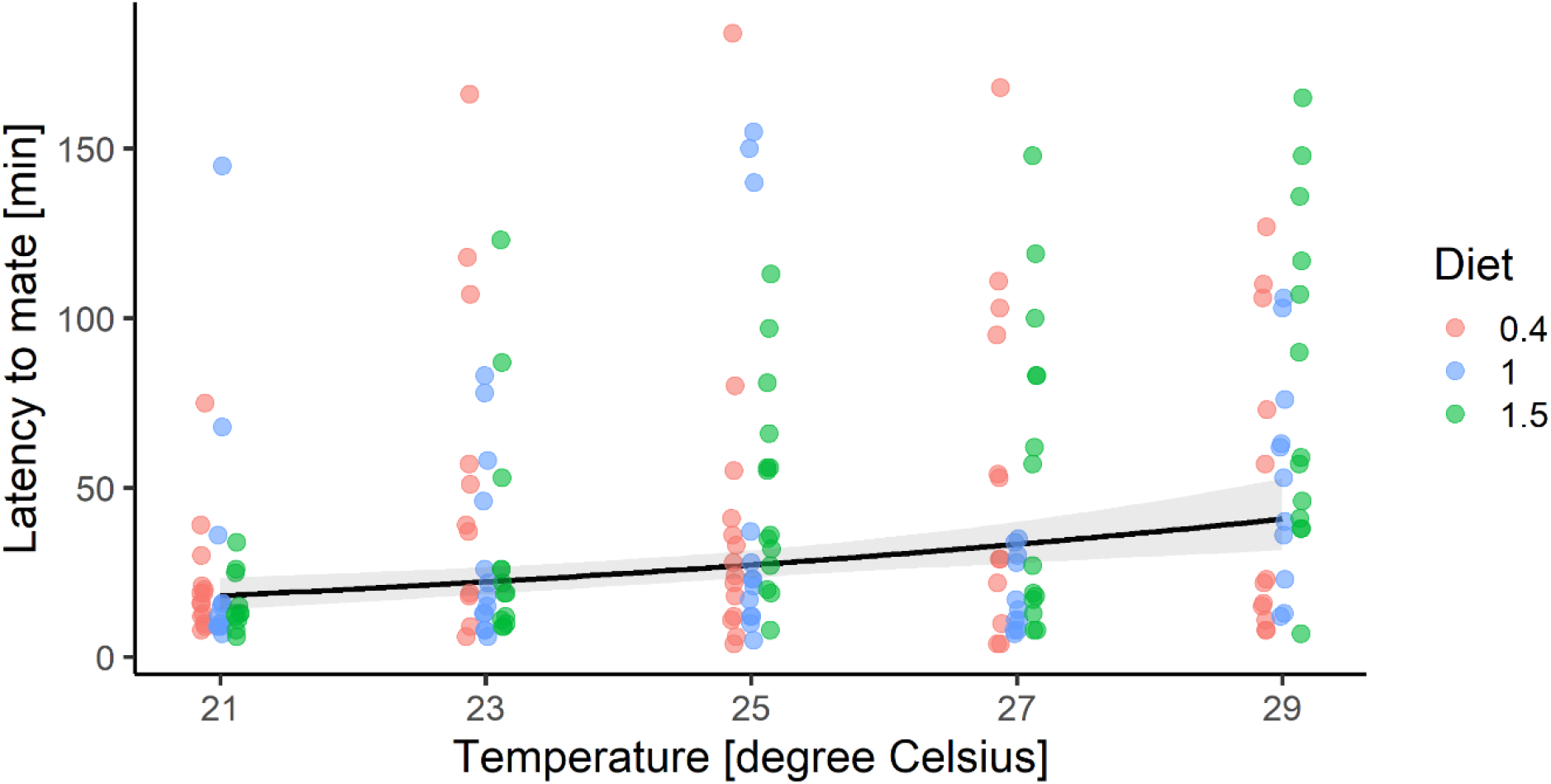
Male latency to mate in generation F0. Shaded areas represent 95% confidence bands.

Maternal temperature and diet treatments did not affect sons’ (F1) latency to mate, mating duration, or mating success (Tables S13, S14, S15). We found no paternal effect on sons’ (F1) latency to mate (Table S16). However, paternal temperature treatment had a non-linear effect on sons’ (F1) mating duration (Figure 5; Table S17; Temperature-dependent smooth model, approximate significance of temperature smooth term: edf = 2.75, F = 3.712, p = 0.049). Paternal diet did not affect sons’ mating performance. The best model of paternal effects on sons’ mating success contained a non-linear temperature term and no diet effect, but the smooth term itself was not significant (Table S15; edf = 1, χ^2^ = 0.003, p = 0.954).

**Figure 5.**
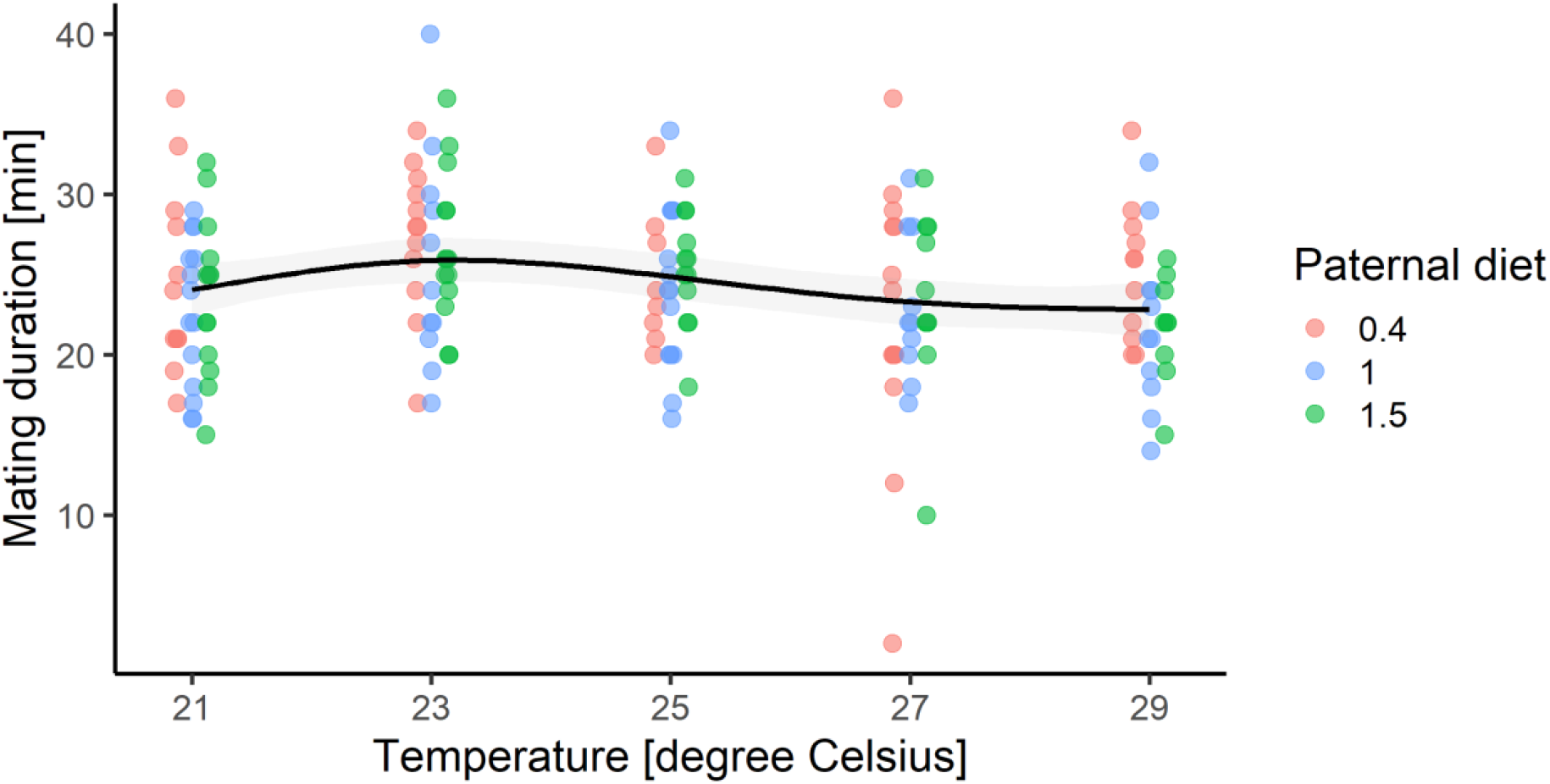
Effects of paternal diet on male mating duration in generation F1. Temperature refers to the experimental temperature fathers of tested males experienced. Shaded areas represent 95% confidence bands.

Grandmaternal temperature had a quadratic effect on grandsons’ (F2) latency to mate, independent of grandmaternal diet (Figure 6; Table S19). Males whose grandmothers had been maintained at 25°C took longer, on average, to start mating compared to males whose grandmothers had been maintained at 21°C or 29°C (Figure 6). Males whose grandfathers had been maintained on low diet had a shorter mating duration compared to males whose grandfathers had been maintained on high diet at 25°C but not at 21 or 29°C (Figure 7; Table S20). For all other mating behaviour traits in F2 and F3 males, we found no evidence of grand-parental or great-grand-parental effects (Tables S21 – S30).

**Figure 6.**
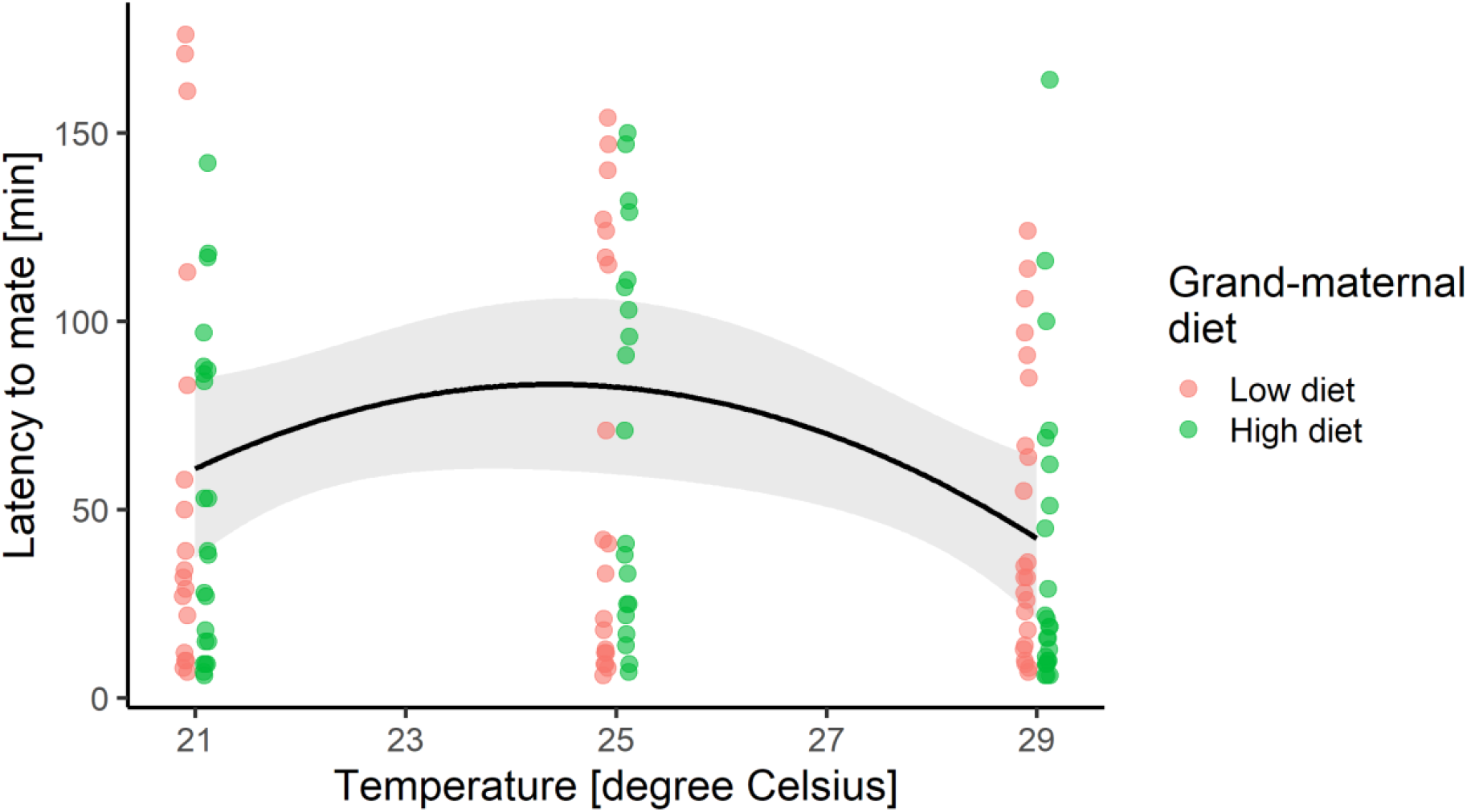
Effects of grand-maternal diet on male latency to mate in generation F2. Temperature refers to the experimental temperature grandmothers of tested males experienced. Shaded areas represent 95% confidence bands.

**Figure 7.**
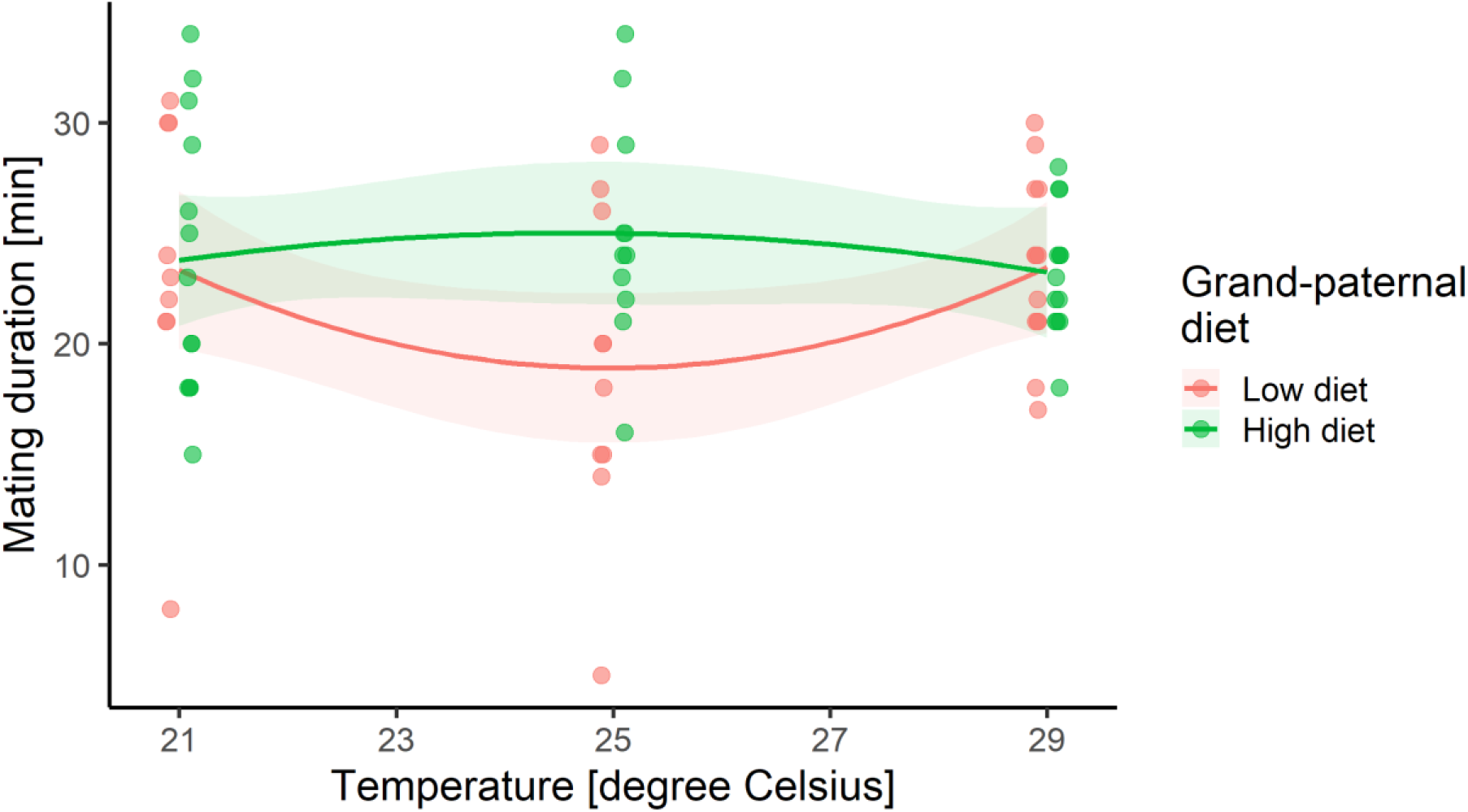
Effects of grand-paternal diet on mating duration in grandsons (generation F2). Temperature refers to the experimental temperature grandfathers of tested males experienced.

## DISCUSSION

We found that reduced protein content in the adult diet increased longevity of *Drosophila melanogaster* females and males when maintained at benign ambient temperature (25-27°C), but not under conditions of cold-stress (21-23°C) or heat-stress (29°C). Indeed, in females, protein restriction resulted in substantially elevated mortality at 21°C and 23°C. In males, the interaction of diet with temperature was less pronounced: protein restriction prolonged life at 25°C and 27°C but had no clear effect on longevity at other temperatures. These results are consistent with previous evidence that dietary restriction increases frailty under some ecologically relevant forms of environmental stress (Burger *et al*. 2006; Adler & Bonduriansky 2014; Savola *et al*. 2021). Our findings suggest that protein-restricted insects are unlikely to achieve extended lifespans in stressful natural environments, and therefore challenge the idea that the longevity-extending effect of DR seen under benign laboratory conditions represents an adaptive survival strategy.

Treatment effects on reproductive performance did not offset the effects of temperature stress on longevity. In females, fecundity of protein-restricted individuals was lower than that of individuals on standard or high-protein diets at all temperatures except 29°C. Interestingly, at cold temperatures (21-23°C), females maintained on high- and standard-protein diets simultaneously maximized both their longevity and their lifetime fecundity, whereas protein-restricted females had dramatically shortened lives and laid fewer eggs over their lifetimes. In males, mating success was not affected by either diet or temperature treatments but male latency to mate (under standard temperature conditions) increased with treatment temperature. Thus, we found no evidence that the failure of protein restriction to prolong life under cold- or hot-temperature stress was associated with positive effects on reproduction. While the costs of reproduction can result in trade-offs between lifespan and reproductive effort (Flatt 2011), our results instead suggest that temperature stress imposes physiological costs that reduce both survival and reproduction. If our results can be generalised to natural populations, they suggest that protein-restricted flies experiencing temperature stress might not only fail to achieve greater longevity but also fail to achieve a higher reproductive rate than fully-fed flies.

Investment in somatic maintenance or fecundity could trade off with investment in offspring quality (Fox & Czesak 2000; Kölliker *et al*. 2015). However, we found no clear evidence that protein-restricted flies produced higher-quality offspring. Low protein maternal diet reduced daughters’ fecundity at both low and high maternal ambient temperatures. Thus, for females experiencing low ambient temperatures, access to abundant dietary protein enhanced longevity, fecundity and offspring quality. Dietary protein could enhance offspring quality by providing the essential building-blocks for yolk synthesis (Mirth *et al*. 2019), and abundant protein could be especially important at low temperatures, which limit flies’ ability to feed (Klepsatel *et al*. 2019). Low protein paternal diet increased daughters’ fecundity, but this effect was only apparent at low paternal temperature treatments (21-23°C). Protein restriction did not enhance F0 male mating performance, nor mating performance of sons (F1), grandsons (F2) or great-grandsons (F3). Indeed, the only effect of diet treatment on mating performance of male descendants was a positive effect of grand-paternal dietary protein at 25°C on mating duration of grandsons (F2). Thus, protein restriction did not induce consistent, positive effects on offspring quality that could offset the negative effects on parental longevity and fecundity.

Rather, the temperature- and diet-induced maternal and paternal effects that we observed were complex and likely to reflect a combination of adaptive and deleterious responses. Environment-induced parental effects could reflect adaptive parental strategies that enhance offspring performance in the environmental conditions that offspring are likely to encounter (Bernardo 1996), or that enhance the performance of offspring produced by parents in high condition (Bonduriansky 2021). Our results suggest that *D. melanogaster* females maximised their condition when ambient temperature was low and dietary protein was abundant, and females maintained under such conditions also appeared to transfer their high condition to their daughters. Alternatively, such effects could occur as deleterious consequences of parental stress (Bernardo 1996; Bell & Hellmann 2019; Bonduriansky 2021). Higher temperature might result in elevated stress for *D. melanogaster* males, potentially explaining why daughters’ fecundity tended to decrease with the ambient temperature experienced by their fathers.

Our results suggest that thermal stress imposes physiological costs that elevate requirements for dietary protein in homeostasis and somatic maintenance. High ambient temperature accelerates metabolic processes in insects, and this probably results in more rapid deterioration of somatic cells and tissues (Mołoń *et al*. 2020). Dietary protein requirements might therefore increase at higher ambient temperatures because of a greater need to repair and replace damaged cells. Moreover, *D. melanogaster* and many other animals respond to heat stress by producing protective heat shock proteins (Tower 2011), and the need to synthesize these proteins could elevate requirements for dietary protein. Because cold temperature slows metabolism in *Drosophila*, the positive effects of dietary protein on survival at colder ambient temperatures are more intriguing. However, some *Drosophila* enzymes exhibit increased activity in response to low temperatures (Burnell *et al*. 1991), and the need to synthesize these enzymes could elevate requirements for protein. Another possibility is that, because cold shock alters concentrations of some amino acids in *Drosophila suzukii* (Enriquez *et al*. 2018), the need to maintain homeostasis under cold conditions could increase dietary requirements for particular amino acids. Studies examining the effects of DR on expression of proteins (e.g. Gao *et al*. 2020), especially if combined with manipulation of ambient temperature, could provide additional clues on how dietary protein requirements for homeostasis and somatic maintenance are altered by thermal stress.

The life-extending effect of DR has been reported in a broad range of animals (Fontana *et al*. 2010), and involves highly conserved physiological pathways (Kapahi *et al*. 2017). This response has therefore been interpreted as a highly conserved physiological mechanism that enhances fitness in insects, mammals and many other animals by helping individuals to survive periods of famine (Kirkwood & Shanley 2005). However, this idea is based on the assumption that dietary restriction tends to prolong life not only under benign laboratory conditions but also under the more stressful conditions experienced by wild animals, including small-bodied animals such as insects. This assumption has rarely been tested, and is challenged by evidence that dietary restriction reduces ability to cope with a range of stresses (Adler & Bonduriansky 2014). Our results show that protein restriction extends lifespan under benign temperature conditions, but fails to extend (and can even shorten) life of flies experiencing thermal stress. Our findings therefore suggest that the lifespan-extending effect of DR (specifically, protein restriction) reported in many laboratory experiments is more plausibly interpreted as an artefact of benign laboratory conditions than as a fitness-enhancing strategy that evolved in natural populations.

However, DR’s effects on frailty appear to depend strongly on the type of stress that experimental animals experience (Mair 2005; Burger *et al*. 2006; Savola *et al*. 2021), and could also be taxon-specific (Adler & Bonduriansky 2014). For example, it is possible that DR could promote extended longevity in large-bodied animals that experience relatively low mortality rates in the wild. Further studies are needed to determine how DR affects longevity in different species, and in response to different types of stress, such as pathogens and parasites, toxins, thermal stress, and interactions between these different types of stress. Further research is also needed to identify the most important sources of stress and mortality in natural populations of insects and other animals. Such work could help to clarify how DR affects survival and fitness in natural populations, how such effects vary across taxa, and how the life-extending effect of DR evolved.

## ACKNOWLEDGEMENTS

We thank Matthew Piper for providing the founders for the Dahomey stock, and undergraduate volunteers Pearl Lei and Sarah Zeigman for counting fly eggs. This research was supported by the Australian Research Council through Discovery Grant DP170102449 to RB.

## Supplementary material

### TABLES

**Table S1.**
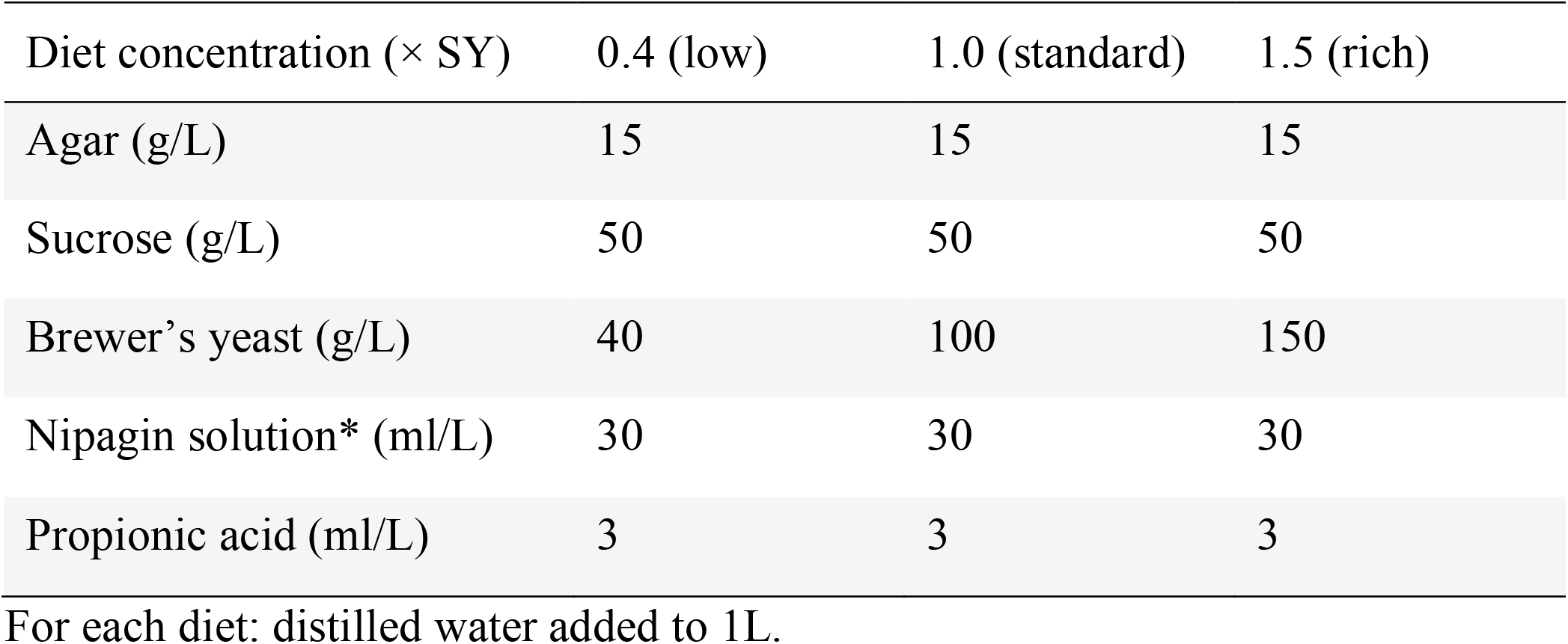
Diet composition.

**Table S2.** Summary statistics for survival of F0 female and male flies.

| Temperature | Diet | mean LS (CI<sub>lower</sub>, CI<sub>upper</sub>; median) |  | N<sub>ind</sub> |
| --- | --- | --- | --- | --- |
|  |  | Females | Males |  |
| 21 | 0.4 | 53.3 (46.7, 60.9; 36.5) | 77.7 (73.0, 82.2; 75) | 100 |
| 21 | 1.0 | 119.0 (114, 123; 126) | 73.6 (69.7, 77.3; 68) | 100 |
| 21 | 1.5 | 122.0 (117, 126; 126) | 77.6 (74.0, 81.5; 76) | 100 |
| 23 | 0.4 | 51.5 (47.5, 55.6; 52) | 48.1 (45.8, 50.3; 47) | 100 |
| 23 | 1.0 | 69.1 (66.6, 70.9; 71.5) | 49.5 (47.3, 51.8; 49) | 100 |
| 23 | 1.5 | 70.1 (68.1, 71.8; 73) | 45.8 (43.8, 47.8; 45) | 100 |
| 25 | 0.4 | 52.4 (49.8, 54.6; 54) | 39.3 (37.8, 40.9; 40) | 100 |
| 25 | 1.0 | 56.4 (54.0, 58.7; 56) | 37.3 (35.8, 38.7; 38) | 100 |
| 25 | 1.5 | 53.9 (51.9, 55.5; 56) | 34.1 (32.7, 35.7; 33) | 100 |
| 27 | 0.4 | 45.1 (43.0, 46.8; 49) | 34.3 (32.8, 35.7; 35) | 100 |
| 27 | 1.0 | 40.4 (38.7, 41.8; 40) | 29.5 (27.9, 30.9; 29.5) | 100 |
| 27 | 1.5 | 40.3 (38.7, 41.6; 42) | 26.1 (25.1, 27.4; 26) | 100 |
| 29 | 0.4 | 38.6 (36.4, 40.4; 40) | 21.3 (19.7, 23; 24) | 100 |
| 29 | 1.0 | 37.7 (35.8, 39.3; 40) | 19.3 (18.1, 20.8; 19) | 100 |
| 29 | 1.5 | 34 (32.2, 35.7; 35) | 20.5 (19.1, 21.8; 24) | 100 |

LS: lifespan, CI: 95% confidence interval, bootstrapped with n = 5000 replicates, with function *groupwiseMean* and option BCa from R package *rcompanion* (). N<sub>ind</sub>: sample size of number of individual flies with available date of death.

**Table S3.** Summary statistics for female fecundity in the F0 generation.

| Temperature | Diet | Mean number of eggs (CI<sub>lower</sub>, CI<sub>upper</sub>; median) | N<sub>vials</sub> |
| --- | --- | --- | --- |
| 21 | 0.4 | 25.7 (17.8, 39.6; 20.5) | 10 |
| 21 | 1.0 | 112.0 (99.3, 125.0; 112) | 10 |
| 21 | 1.5 | 207.0 (192.0, 219.0; 207) | 10 |
| 23 | 0.4 | 30.9 (21.5, 40.4; 27.5) | 10 |
| 23 | 1.0 | 134.0 (114.0, 174.0; 121) | 10 |
| 23 | 1.5 | 203.0 (174.0, 234.0; 208) | 10 |
| 25 | 0.4 | 30.1 (20.7, 56.9; 21.5) | 10 |
| 25 | 1.0 | 80.8 (58.6, 114.0; 62) | 10 |
| 25 | 1.5 | 132.0 (104.0, 155.0; 141) | 10 |
| 27 | 0.4 | 26.0 (14.8, 51.5; 17) | 10 |
| 27 | 1.0 | 123.0 (89.4, 159.0; 124) | 10 |
| 27 | 1.5 | 71.5 (50.0, 111.0; 69) | 10 |
| 29 | 0.4 | 31.2 (21.4, 47.3; 23.5) | 10 |
| 29 | 1.0 | 34.6 (23.2, 46.6; 30.5) | 10 |
| 29 | 1.5 | 25.4 (12.0, 49.7; 10.5) | 10 |

Mean number of eggs of the total number of eggs per vial, which was calculated as the sum of eggs over 5 weeks for each vial (see Methods), CI: 95% confidence interval, bootstrapped with $n = 5000$ replicates, with function *groupwiseMean* and option BCa from R package rcompanion (). N<sub>ind</sub>: sample size of number of individual flies with available date of death.

**Table S4.** Cox GAM survival model comparisons for generation F0.

| Sex | Model | df | AIC | ΔAIC |
| --- | --- | --- | --- | --- |
| Female | **Diet-specific smooths** | 64 | 17643.21 | 0.00 |
|  | Diet-specific intercept | 107 | 17686.36 | 43.15 |
|  | No diet effect | 109 | 17689.33 | 46.11 |
| Male | **Diet-specific intercept** | 84 | 17295.81 | 0.00 |
|  | No diet effect | 92 | 17302.19 | 6.37 |
|  | Diet-specific smooths | 94 | 17304.73 | 8.92 |

**Table S5.** Pairwise logrank survival model comparisons (corrected for multiple comparisons) for generation F0.

| Sex | Comparison | Temperature |  |  |  |  |
| --- | --- | --- | --- | --- | --- | --- |
|  |  | 21 | 23 | 25 | 27 | 29 |
| Female | 0.4 vs. 1.0 | **< 0.001** | **< 0.001** | **0.019** | **< 0.001** | **0.016** |
|  | 0.4 vs. 1.5 | **< 0.001** | **< 0.001** | 0.719 | **< 0.001** | **< 0.001** |
|  | 1.0 vs. 1.5 | 0.123 | 0.486 | **0.019** | 0.514 | **0.007** |
| Male | 0.4 vs. 1.0 | 0.248 | 0.399 | **0.037** | **< 0.001** | 0.061 |
|  | 0.4 vs. 1.5 | 0.428 | 0.073 | **< 0.001** | **< 0.001** | 0.212 |
|  | 1.0 vs. 1.5 | 0.352 | **0.024** | **0.018** | **< 0.001** | 0.146 |

**Table S6.** Cox survival model results for female offspring (generation F1).

| Model term | coef | exp(coef) | se(coef) | z | p |
| --- | --- | --- | --- | --- | --- |
| Parental sex | 0.361 | 1.435 | 1.056 | 0.34 | 0.733 |
| Parental temperature | 0.052 | 1.054 | 0.034 | 1.53 | 0.127 |
| Parental diet (Standard) | −0.813 | 0.444 | 1.220 | −0.67 | 0.505 |
| Parental diet (Rich) | 1.096 | 2.992 | 1.221 | 0.90 | 0.369 |
| Parental sex × Parental temperature | −0.025 | 0.975 | 0.042 | −0.61 | 0.545 |
| Parental sex × Parental diet (Standard) | 0.738 | 2.092 | 1.475 | 0.50 | 0.617 |
| Parental sex × Parental diet (Rich) | 0.120 | 1.127 | 1.480 | 0.08 | 0.935 |
| Parental temperature × Parental diet<br>(Standard) | 0.018 | 1.019 | 0.048 | 0.38 | 0.705 |
| Parental temperature × Parental diet<br>(Rich) | −0.059 | 0.942 | 0.049 | −1.22 | 0.222 |
| Parental sex × Parental temperature ×<br>Parental diet (Standard) | −0.016 | 0.984 | 0.059 | −0.28 | 0.778 |
| Parental sex × Parental temperature ×<br>Parental diet (Rich) | 0.002 | 1.002 | 0.059 | 0.04 | 0.967 |

**Table S7.** Cox survival model results for male offspring (generation F1).

| Model term | coef | exp(coef) | se(coef) | z | p |
| --- | --- | --- | --- | --- | --- |
| Parental sex | −1.250 | 0.286 | 1.014 | −1.23 | 0.218 |
| Parental temperature | −0.004 | 0.996 | 0.030 | −0.14 | 0.891 |
| Parental diet (Standard) | −0.172 | 0.842 | 1.056 | −0.16 | 0.871 |
| Parental diet (Rich) | −0.688 | 0.502 | 1.069 | −0.64 | 0.520 |
| Parental sex × Parental temperature | 0.042 | 1.043 | 0.040 | 1.05 | 0.294 |
| Parental sex × Parental diet (Standard) | 1.852 | 6.376 | 1.449 | 1.28 | 0.201 |
| Parental sex × Parental diet (Rich) | 0.256 | 1.291 | 1.454 | 0.18 | 0.860 |
| Parental temperature × Parental diet<br>(Standard) | 0.004 | 1.004 | 0.042 | 0.10 | 0.917 |
| Parental temperature × Parental diet<br>(Rich) | 0.023 | 1.023 | 0.043 | 0.54 | 0.591 |
| Parental sex × Parental temperature ×<br>Parental diet (Standard) | −0.070 | 0.932 | 0.058 | −1.22 | 0.225 |
| Parental sex × Parental temperature ×<br>Parental diet (Rich) | <0.001 | 1.001 | 0.058 | 0.01 | 0.992 |

**Table S8.**
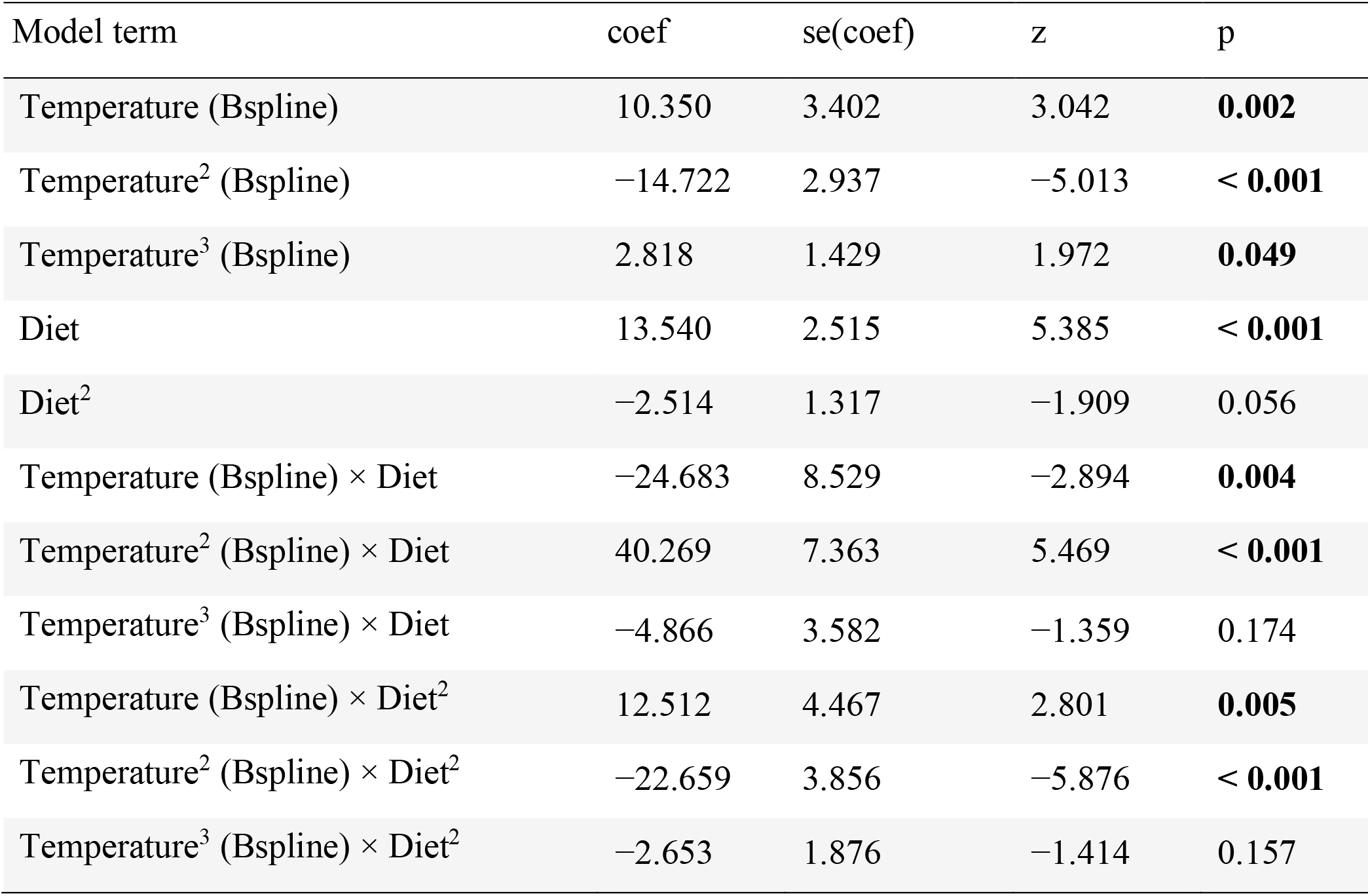
Robust GLM model results for female fecundity (generation F0).

**Table S9.** Robust GLM model results for female fecundity (generation F1).

| Model term | coef | se(coef) | z | p |
| --- | --- | --- | --- | --- |
| Parental sex | 7.791 | 3.172 | 2.457 | **0.014** |
| Parental temperature (Bspline) | -9.880 | 7.607 | -1.299 | 0.194 |
| Parental temperature<sup>2</sup> (Bspline) | 18.031 | 6.567 | 2.746 | **0.006** |
| Parental temperature<sup>3</sup> (Bspline) | -6.398 | 3.194 | -2.003 | **0.045** |
| Parental diet | -3.041 | 5.623 | -0.541 | 0.589 |
| Parental diet<sup>2</sup> | 4.427 | 2.945 | 1.503 | 0.133 |
| Parental diet × Parental sex | -4.162 | 7.952 | -0.523 | 0.601 |
| Parental sex × Parental temperature (Bspline) | 16.466 | 10.757 | 1.531 | 0.126 |
| Parental sex × Parental temperature<sup>2</sup> (Bspline) | -34.570 | 9.287 | -3.722 | **< 0.001** |
| Parental sex × Parental temperature<sup>3</sup> (Bspline) | -1.759 | 4.518 | -0.389 | 0.697 |
| Parental diet × Parental temperature (Bspline) | 32.829 | 19.071 | 1.721 | 0.085 |
| Parental diet × Parental temperature<sup>2</sup> (Bspline) | -37.542 | 16.464 | -2.280 | **0.023** |
| Parental diet × Parental temperature<sup>3</sup> (Bspline) | 21.796 | 8.009 | 2.721 | **0.007** |
| Parental diet<sup>2</sup> × Parental sex | -2.673 | 4.164 | -0.642 | 0.521 |
| Parental diet<sup>2</sup> × Parental temperature (Bspline) | -19.446 | 9.988 | -1.947 | 0.052 |
| Parental diet<sup>2</sup> × Parental temperature<sup>2</sup> (Bspline) | 13.185 | 8.622 | 1.529 | 0.126 |
| Parental diet<sup>2</sup> × Parental temperature<sup>3</sup> (Bspline) | -13.393 | 4.194 | -3.193 | **0.001** |
| Parental sex × Parental diet × Temperature (Bspline) | -66.497 | 26.971 | -2.466 | **0.014** |
| Parental sex × Parental diet × Temperature<sup>2</sup><br>(Bspline) | 87.122 | 23.284 | 3.742 | **< 0.001** |
| Parental sex × Parental diet × Temperature<sup>3</sup><br>(Bspline) | -11.416 | 11.327 | -1.008 | 0.314 |
| Parental sex × Parental diet<sup>2</sup> × Temperature<br>(Bspline) | 39.804 | 14.125 | 2.818 | **0.005** |
| Parental sex × Parental diet<sup>2</sup> × Temperature<sup>2</sup><br>(Bspline) | -40.379 | 12.193 | -3.312 | **< 0.001** |
| Parental sex × Parental diet<sup>2</sup> × Temperature<sup>3</sup><br>(Bspline) | 9.538 | 5.932 | 1.608 | 0.108 |

**Table S10.** GAMM male latency to mate model comparison for generation F0. Columns ‘Temperature’ and ‘Diet’ indicate whether these predictors (either as smooths or as linear/intercepts) were part of the specific model.

| Model | Temperature | Diet | df | AIC | ΔAIC |
| --- | --- | --- | --- | --- | --- |
| **Temperature-dependent smooth** | x |  | 5 | 517.3 | 0.00 |
| **Linear temperature** | x |  | 4 | 519.0 | 1.69 |
| Temperature-dependent smooth and linear diet | x | x | 7 | 522.0 | 4.74 |
| Mean only (Null model) |  |  | 3 | 523.4 | 6.11 |
| Temperature-dependent and diet-specific smooths | x | x | 11 | 526.2 | 8.94 |

**Table S11.** GAMM male mating duration model comparison for generation F0. Columns ‘Temperature’ and ‘Diet’ indicate whether these predictors (either as smooths or as linear/intercepts) were part of the specific model.

| Model | Temperature | Diet | df | AIC | ΔAIC |
| --- | --- | --- | --- | --- | --- |
| **Mean only (Null model)** |  |  | 3 | 1109.5 | 0.00 |
| **Temperature-dependent and diet-specific smooths** | x | x | 11 | 1110.0 | 0.49 |
| **Temperature-dependent smooth** | x |  | 5 | 1111.5 | 1.94 |
| Temperature-dependent smooth and linear diet | x | x | 7 | 1111.8 | 2.28 |
| Linear temperature | x |  | 4 | 1113.8 | 4.24 |

**Table S12.** GAMM male mating success model comparison for generation F0. Columns ‘Temperature’ and ‘Diet’ indicate whether these predictors (either as smooths or as linear/intercept) were part of the specific model.

| Model | Temperature | Diet | df | AIC | ΔAIC |
| --- | --- | --- | --- | --- | --- |
| **Mean only (Null model)** |  |  | 3 | 71.6 | 0.00 |
| Temperature-dependent smooth | x |  | 5 | 75.6 | 3.99 |
| Temperature-dependent and diet-specific smooths | x | x | 11 | 91.5 | 19.86 |
| Linear temperature | x |  | 4 | 144.1 | 72.48 |
| Temperature-dependent smooth and linear diet | x | x | 7 | 148.8 | 77.17 |

**Table S13.** Maternal effect GAMM male latency to mate model comparison for generation F1. Columns ‘Temperature’ and ‘Diet’ indicate whether these predictors (either as smooths or as linear/intercept term) were part of the specific model.

| Model | Temperature | Diet | df | AIC | ΔAIC |
| --- | --- | --- | --- | --- | --- |
| **Mean only (Null model)** |  |  | 3 | 460.1 | 0.00 |
| Temperature-dependent smooth | x |  | 5 | 465.0 | 4.89 |
| Linear temperature | x |  | 4 | 466.7 | 6.58 |
| Temperature-dependent smooth and linear diet | x | x | 7 | 470.0 | 9.86 |
| Temperature-dependent and diet-specific smooths | x | x | 11 | 1625.7 | 1165.52 |

**Table S14.** Maternal effect GAMM male mating duration model comparison for generation F1. Columns ‘Temperature’ and ‘Diet’ indicate whether these predictors (either as smooths or as linear/intercept term) were part of the specific model.

| Model | Temperature | Diet | df | AIC | ΔAIC |
| --- | --- | --- | --- | --- | --- |
| **Temperature-dependent and diet-specific smooths** | x | x | 11 | 1024.3 | 0.00 |
| **Mean only (Null model)** |  |  | 3 | 1025.6 | 1.34 |
| Temperature-dependent smooth and linear diet | x | x | 7 | 1027.2 | 2.93 |
| Temperature-dependent smooth | x |  | 5 | 1027.6 | 3.35 |
| Linear temperature | x |  | 4 | 1029.3 | 5.04 |

**Table S15.** Maternal effect GAMM male mating success model comparison for generation F1. Columns ‘Temperature’ and ‘Diet’ indicate whether these predictors (either as smooths or as linear/intercept term) were part of the specific model.

| Model | Temperature | Diet | df | AIC | ΔAIC |
| --- | --- | --- | --- | --- | --- |
| **Mean only (Null model)** |  |  | 3 | 239.6 | 0.00 |
| **Temperature-dependent and diet-specific smooths** | x | x | 11 | 241.0 | 1.38 |
| Linear temperature | x |  | 4 | 241.6 | 2.00 |
| Temperature-dependent smooth and linear diet | x | x | 7 | 242.2 | 2.62 |
| Temperature-dependent smooth | x |  | 5 | 243.6 | 4.00 |

**Table S16.** Paternal effect GAMM male latency to mate model comparison for generation F1. Columns ‘Temperature’ and ‘Diet’ indicate whether these predictors (either as smooths or as linear/intercept term) were part of the specific model.

| Model | Temperature | Diet | df | AIC | ΔAIC |
| --- | --- | --- | --- | --- | --- |
| **Mean only (Null model)** |  |  | 3 | 513.2 | 0.00 |
| Temperature-dependent smooth | x |  | 5 | 516.7 | 3.50 |
| Linear temperature | x |  | 4 | 518.4 | 5.20 |
| Temperature-dependent smooth and linear diet | x | x | 7 | 523.4 | 10.21 |
| Temperature-dependent and diet-specific smooths | x | x | 11 | 530.6 | 17.37 |

**Table S17.** Paternal effect GAMM male mating duration model comparison for generation F1. Columns ‘Temperature’ and ‘Diet’ indicate whether these predictors (either as smooths or as linear/intercept term) were part of the specific model.

| Model | Temperature | Diet | df | AIC | ΔAIC |
| --- | --- | --- | --- | --- | --- |
| **Temperature-dependent smooth and linear diet** | x | x | 7 | 1074.1 | 0.00 |
| **Temperature-dependent smooth** | x |  | 5 | 1075.8 | 1.69 |
| Temperature-dependent and diet-specific smooths | x | x | 11 | 1076.8 | 2.67 |
| Mean only (Null model) |  |  | 3 | 1078.7 | 4.59 |
| Linear temperature | x |  | 4 | 1080.0 | 5.92 |

**Table S18.** Paternal effect GAMM male mating success model comparison for generation F1. Columns ‘Temperature’ and ‘Diet’ indicate whether these predictors (either as smooths or as linear/intercept term) were part of the specific model.

| Model | Temperature | Diet | df | AIC | ΔAIC |
| --- | --- | --- | --- | --- | --- |
| **Temperature-dependent smooth** | x |  | 5 | 123.8 | 0.00 |
| Temperature-dependent smooth and linear diet | x | x | 7 | 127.7 | 3.96 |
| Temperature-dependent and diet-specific smooths | x | x | 11 | 139.2 | 15.45 |
| Linear temperature | x |  | 4 | 198.0 | 74.25 |
| Mean only (Null model) |  |  | 3 | 200.8 | 76.99 |

**Table S19.** Final grand-maternal effect LMM of male latency to mate for generation F2.

| Model term | coef | se(coef) | df | t | p |
| --- | --- | --- | --- | --- | --- |
| Grandparental temperature | 94.242 | 44.083 | 56 | 2.138 | **0.037** |
| Grandparental temperature<sup>2</sup> | −1.931 | 0.879 | 56 | −2.198 | **0.032** |

**Table S20.** Final grand-paternal effect LMM of mating duration of grandsons (generation F2).

| Model term | coef | se(coef) | df | t | p |
| --- | --- | --- | --- | --- | --- |
| Grandparental temperature | −14.039 | 6.577 | 63.00 | −2.135 | **0.037** |
| Grandparental diet (1.5) | −226.092 | 111.122 | 63.00 | −2.035 | **0.046** |
| Grandparental temperature<sup>2</sup> | 0.281 | 0.131 | 63.00 | 2.149 | **0.036** |
| Grandparental temperature × Grandparental diet (1.5) | 18.659 | 8.998 | 63.00 | 2.074 | **0.042** |
| Grandparental diet (1.5) × Grandparental temperature<sup>2</sup> | −0.375 | 0.179 | 63.00 | −2.090 | **0.041** |

**Table S21.**
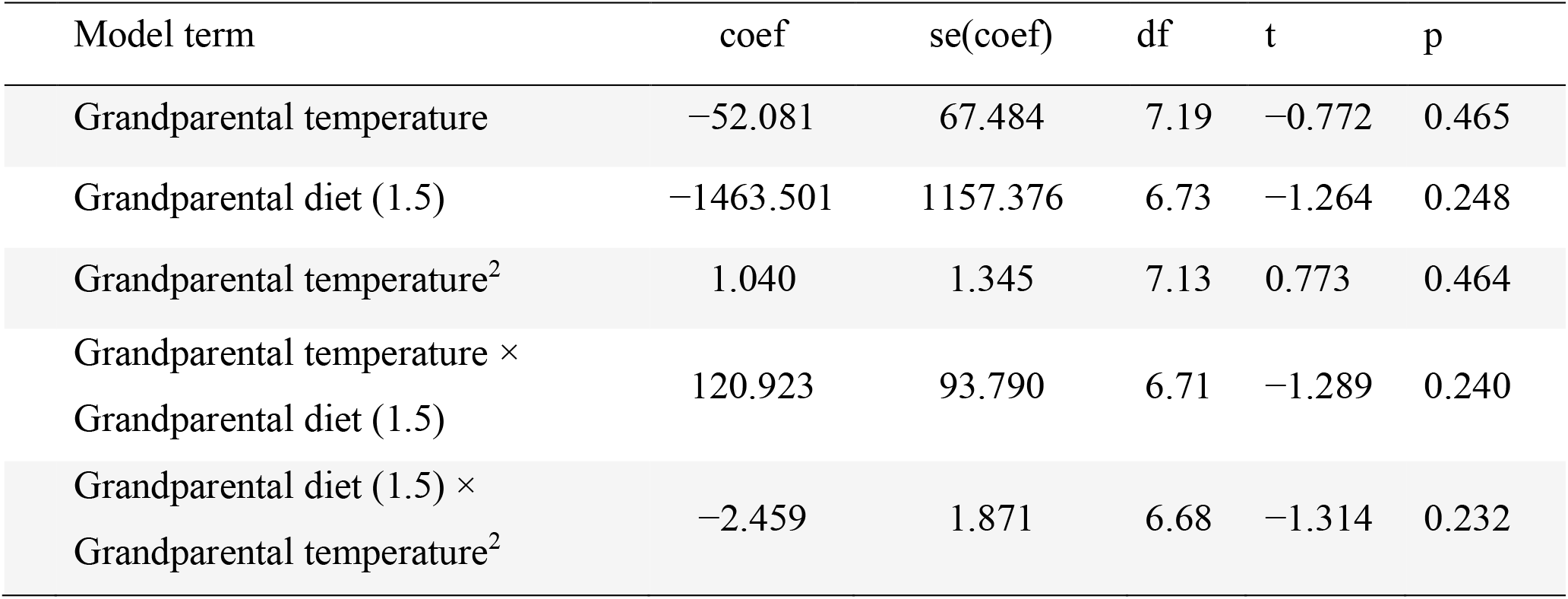
Global grand-paternal effect LMM of male latency to mate for generation F2.

**Table S22.** Global grand-maternal effect LMM of mating duration of grandsons (generation F2).

| Model term | coef | se(coef) | df | t | p |
| --- | --- | --- | --- | --- | --- |
| Grandparental temperature | 3.698 | 6.253 | 5.47 | 0.591 | 0.578 |
| Grandparental diet (1.5) | −80.045 | 110.251 | 6.20 | −0.726 | 0.494 |
| Grandparental temperature<sup>2</sup> | −0.064 | 0.125 | 5.43 | −0.513 | 0.682 |
| Grandparental temperature ×<br>Grandparental diet (1.5) | 7.00 | 8.925 | 6.13 | 0.785 | 0.462 |
| Grandparental diet (1.5) ×<br>Grandparental temperature<sup>2</sup> | −0.148 | 0.178 | 6.08 | −0.829 | 0.438 |

**Table S23.** Global grand-maternal effect GLM of mating success of grandsons (generation F2).

| Model term | coef | se(coef) | t | p |
| --- | --- | --- | --- | --- |
| Grandparental temperature | 0.554 | 2.327 | 0.238 | 0.812 |
| Grandparental diet (1.5) | 26.867 | 40.571 | 0.662 | 0.510 |
| Grandparental temperature<sup>2</sup> | −0.010 | 0.047 | −0.220 | 0.826 |
| Grandparental temperature ×<br>Grandparental diet (1.5) | −2.285 | 3.307 | −0.691 | 0.492 |
| Grandparental diet (1.5) ×<br>Grandparental temperature<sup>2</sup> | 0.048 | 0.066 | 0.721 | 0.473 |

**Table S24.**
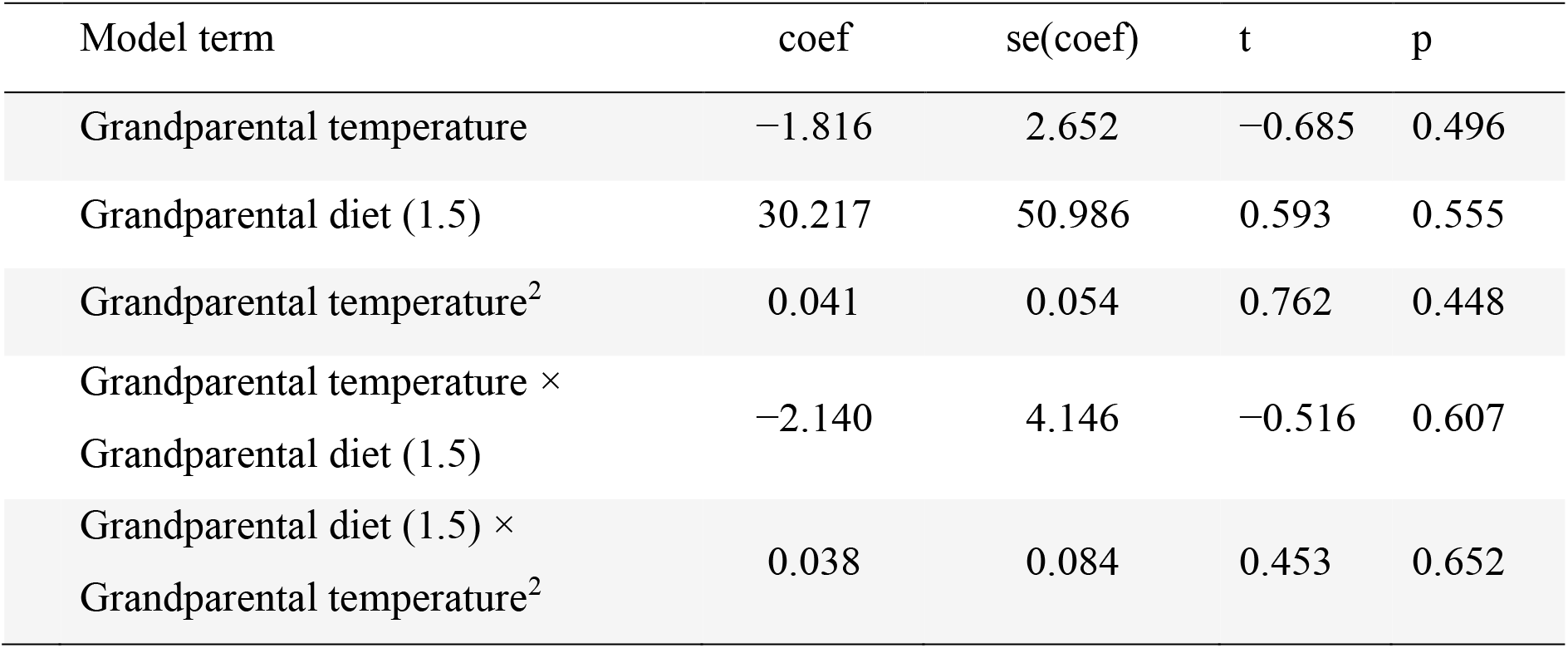
Global grand-paternal effect GLM of mating success of grandsons (generation F2).

| Model term | coef | se(coef) | t | p |
| --- | --- | --- | --- | --- |
| Grandparental temperature | −1.816 | 2.652 | −0.685 | 0.496 |
| Grandparental diet (1.5) | 30.217 | 50.986 | 0.593 | 0.555 |
| Grandparental temperature<sup>2</sup> | 0.041 | 0.054 | 0.762 | 0.448 |
| Grandparental temperature ×<br>Grandparental diet (1.5) | −2.140 | 4.146 | −0.516 | 0.607 |
| Grandparental diet (1.5) ×<br>Grandparental temperature<sup>2</sup> | 0.038 | 0.084 | 0.453 | 0.652 |

**Table S25.** Global great-grand-maternal effect LMM of latency to mate of great-grandsons (generation F3).

| Model term | coef | se(coef) | df | t | p |
| --- | --- | --- | --- | --- | --- |
| Great-grandparental temperature | 16.535 | 43.194 | 6.33 | 0.383 | 0.714 |
| Great-grandparental diet (1.5) | −196.634 | 760.138 | 6.51 | −0.259 | 0.804 |
| Great-grandparental temperature<sup>2</sup> | −0.385 | 0.861 | 6.28 | −0.448 | 0.670 |
| Great-grandparental temperature ×<br>Great-grandparental diet (1.5) | 13.615 | 61.637 | 6.50 | 0.221 | 0.832 |
| Great-grandparental diet (1.5) ×<br>Great-grandparental temperature<sup>2</sup> | −0.227 | 1.230 | 6.48 | −0.185 | 0.859 |

**Table S26.**
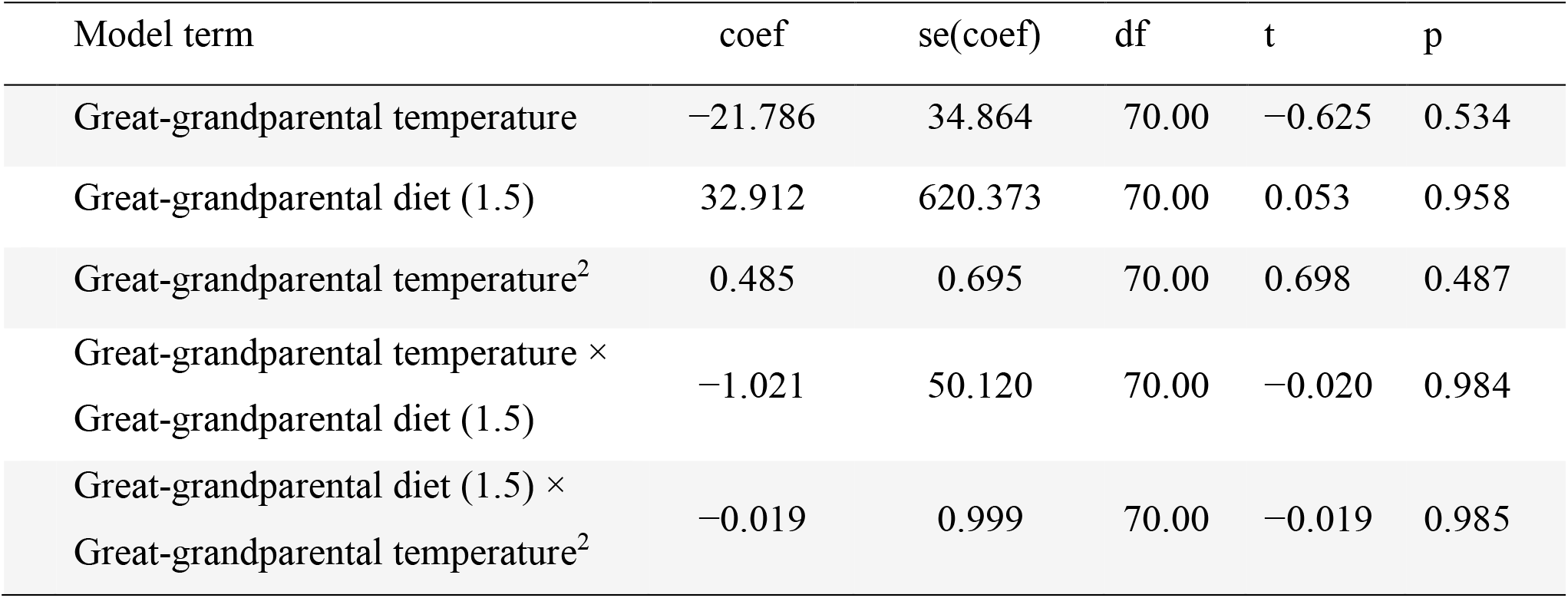
Global great-grand-paternal effect LMM of latency to mate of great-grandsons (generation F3).

**Table S27.** Global great-grand-maternal effect LMM of mating duration of great-grandsons (generation F3).

| Model term | coef | se(coef) | df | t | p |
| --- | --- | --- | --- | --- | --- |
| Great-grandparental temperature | −4.372 | 9.317 | 66.00 | −0.469 | 0.640 |
| Great-grandparental diet (1.5) | −101.519 | 164.322 | 66.00 | −0.618 | 0.539 |
| Great-grandparental temperature<sup>2</sup> | 0.099 | 0.186 | 66.00 | 0.536 | 0.594 |
| Great-grandparental temperature ×<br>Great-grandparental diet (1.5) | 8.872 | 13.323 | 66.00 | 0.666 | 0.508 |
| Great-grandparental diet (1.5) ×<br>Great-grandparental temperature<sup>2</sup> | −0.191 | 0.266 | 66.00 | −0.717 | 0.476 |

**Table S28.** Global great-grand-paternal effect LMM of mating duration of great-grandsons (generation F3).

| Model term | coef | se(coef) | df | t | p |
| --- | --- | --- | --- | --- | --- |
| Great-grandparental temperature | −4.336 | 3.435 | 5.95 | −1.262 | 0.254 |
| Great-grandparental diet (1.5) | −77.546 | 61.064 | 6.40 | −1.270 | 0.248 |
| Great-grandparental temperature<sup>2</sup> | 0.085 | 0.068 | 5.909 | 1.240 | 0.262 |
| Great-grandparental temperature ×<br>Great-grandparental diet (1.5) | 6.509 | 4.934 | 6.32 | 1.319 | 0.233 |
| Great-grandparental diet (1.5) ×<br>Great-grandparental temperature<sup>2</sup> | −0.135 | 0.098 | 6.26 | −1.375 | 0.216 |

**Table S29.**
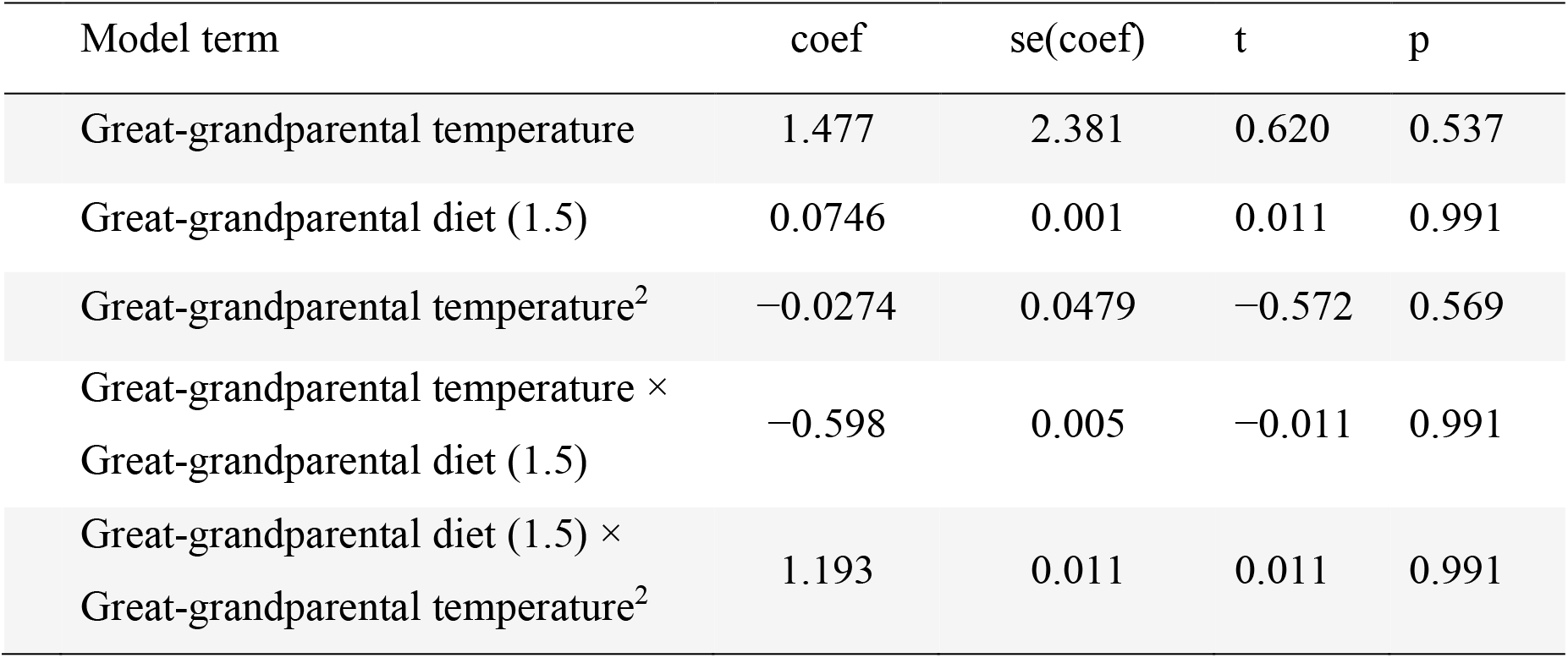
Global great-grand-maternal effect GLM of mating success of great-grandsons (generation F3).

**Table S30.**
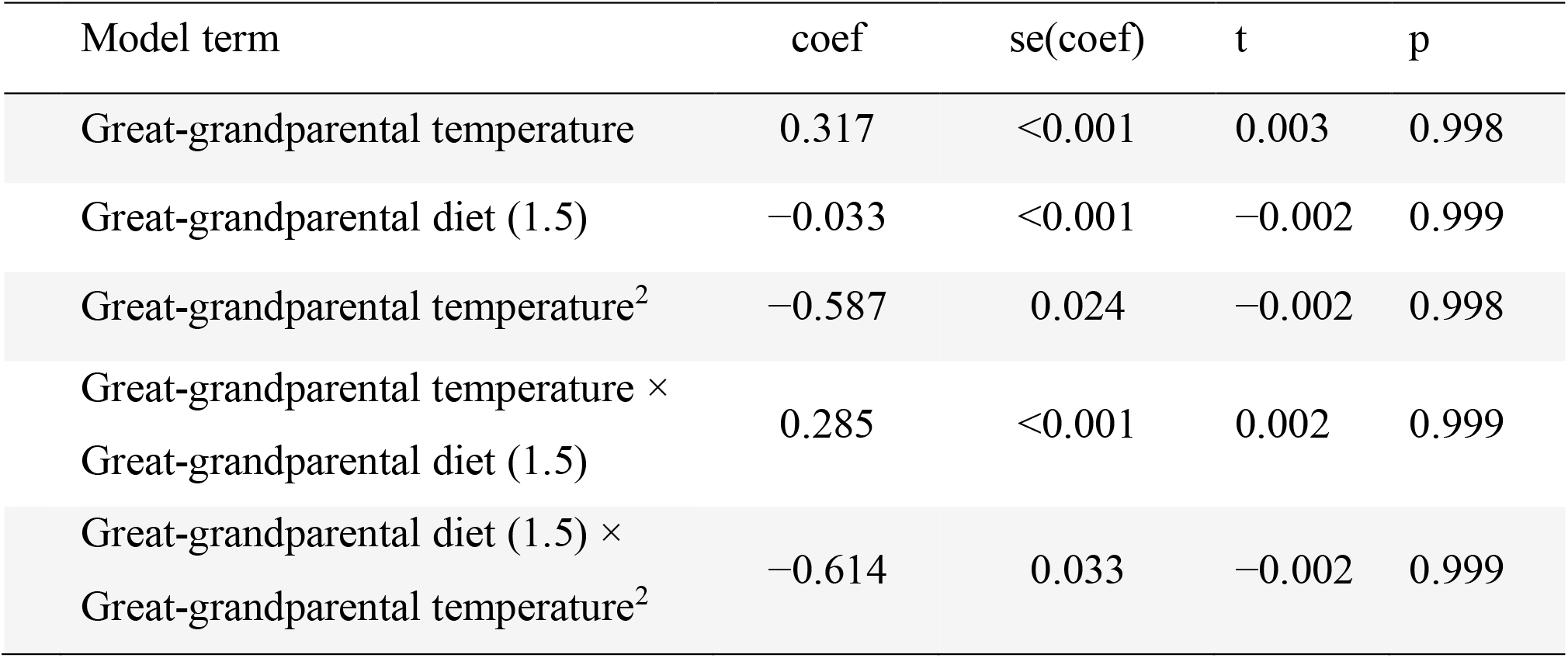
Global great-grand-paternal effect GLM of mating success of great-grandsons (generation F3).

